# Methyltransferases in Candidate Phyla Radiation: A Weapon or a Simple Shield?

**DOI:** 10.64898/2026.07.13.738296

**Authors:** Riyad Razzouk, Linda Hadjadj, Muriel Militello, Mamadou Beye, Lucile Pinault, Jean-Marc Rolain, Fadi Bittar

## Abstract

The Candidate Phyla Radiation (CPR) consists of microorganisms with highly reduced genomes and limited metabolic capabilities, including the inability to synthesize nucleotides and amino acids. Previous studies revealed an unexpected abundance of antimicrobial resistance-related genes in CPR genomes, with methyltransferases (MTs) accounting for up to 47.34% of detected resistance determinants. Here, we investigated the distribution, diversity, and potential functions of MTs across 12,552 CPR genomes, focusing on their roles in antimicrobial resistance and restriction–modification (R–M) systems. Sequence similarity searches were performed using BLASTp, followed by domain validation with the NCBI Conserved Domain Database (CDD). Putative restriction endonucleases (REs) associated with these systems were also identified and analyzed. We detected 20,232 resistance-associated MTs, 7,391 hsdM-like MTs, and 4,737 REs, revealing a high prevalence of putatively functional MTs relative to REs across CPR genomes. Functional CPR MTs and REs displayed extensive sequence divergence from known bacterial homologs and exhibited greater domain diversity and functional complexity than their bacterial counterparts. Selected high-confidence MT candidates were heterologously expressed in *Escherichia coli* BL21 and characterized using BIOLOG phenotypic assays and antimicrobial susceptibility testing. Transformed strains showed enhanced utilization of methyl pyruvate as a carbon source, whereas no significant antimicrobial resistance phenotype was detected.

Collectively, these findings suggest that CPR-encoded MTs are primarily associated with R–M systems, supporting defense against foreign DNA and facilitating nucleotide scavenging rather than direct antimicrobial resistance. This study provides new insights into the adaptive strategies of CPR bacteria and the multifunctional roles of MTs in metabolically streamlined microorganisms.

**Importance:** Candidate Phyla Radiation (CPR) microorganisms represent a major fraction of Earth’s microbial diversity, yet their biology remains poorly understood. We show that CPR genomes contain large numbers of methyltransferases and associated restriction enzymes, suggesting the existence of a specialized DNA-processing system. Because CPR organisms have highly reduced metabolic capabilities, this system may help them recover valuable nucleotides from external DNA while simultaneously protecting their genomes from foreign genetic material. We also found that CPR enzymes are highly distinct from those described in other bacteria and display remarkable functional diversity, highlighting the unique evolutionary trajectories of these microorganisms. By providing the first large-scale characterization of these enzymes in CPR, our study offers new insights into how these widespread microbes survive, acquire resources, and interact with their environment, expanding our understanding of microbial diversity and adaptation.

## Introduction

Candidate Phyla Radiation (CPR) microbes have recently been established as a monophyletic group distinct from previously recognized bacterial domains, based on rhizomal and phylogenetic reclassifications (1–4). These microorganisms are highly ubiquitous (5) and have been detected across a wide range of environments, including soils (6), subsurface ecosystems (7), marine sediments (8), dolphin oral cavities (9), drinking water systems (10), and human tissues (11).

CPR microbes are estimated to account for approximately 26% of microbial diversity (12). They are characterized by extremely small cell sizes (approximately 200–300 nm) and highly reduced genomes, typically smaller than 1 Mb (12,13). Although many CPR organisms exhibit parasitic lifestyles (14), some studies classify them as episymbionts. Regardless of lifestyle classification, CPRs maintain close interactions with their bacterial hosts, mediated in part by type IV pilus systems that facilitate cell–cell adhesion, twitching motility, and uptake of specific metabolites (15,16). Additional communication mechanisms, including quorum-sensing–related repertoires, have also been described (14).

A defining feature of CPR genomes is the widespread absence of genetic components required for complete metabolic pathways (3). In particular, the lack of tricarboxylic acid (TCA) cycle and respiratory pathway genes has led to the classification of many CPR lineages as fermentative organisms, as first demonstrated in complete Peregrinibacteria genomes (17). Across CPR phyla, the inability to synthesize nucleotides and amino acids is consistently observed (15), reinforcing their dependence on host-derived metabolic resources.

In parallel, large-scale genomic analyses have revealed a surprisingly diverse repertoire of antibiotic resistance–like genes encoded by CPR genomes (12). These findings raise important questions regarding the evolutionary origins of resistance determinants that predate the anthropogenic use of antibiotics (18). Indeed, resistance-associated genes have been detected in metagenomic DNA from microbial communities dating back to more than 5,000 years (18), including in ancient dental calculus samples. Separately, members of the Candidate Phyla Radiation (CPR) have been identified in human-associated microbiomes, including Neanderthal dental calculus (19), making resistance gene analyses more significant. Members of the Saccharibacteria phylum have been detected at high frequency in mammalian tissues, particularly the TM7-G1 lineage, with TM7-G3 and TM7-G6 detected at lower frequencies, alongside representatives of “Candidatus Absconditabacteria”. In addition, members of the Gracilibacteria phylum have also been identified in human oral cavities (19,20).

Following the identification of putative resistance proteins in CPR genomes (12), attention shifted toward CPR-encoded beta-lactamases (21). These enzymes were shown to possess functional properties extending beyond antibiotic degradation, including RNase activity indicative of broader hydrolytic capabilities. Subsequent work by Maatouk et al. (2023) examined CPR metallo-beta-lactamases in comparison with bacterial counterparts, revealing an expanded functional repertoire (22). As a result, CPR beta-lactamase-like proteins were reclassified as metallo-hydrolases, highlighting the multifunctional nature of CPR-encoded enzymes.

Among resistance-like proteins identified in CPR genomes, methyltransferases were particularly abundant, accounting for 47.34% of total detected hits (12). Given their known roles in antimicrobial resistance, epigenetic regulation, and protection of self-DNA within restriction– modification systems (23), methyltransferases represent strong candidates for multifunctional roles in CPR biology. Both restriction (*hsdR*) and methylation genes (*hsdM*) are referred to as host specificity determinants (hsd) (42). Accordingly, the present study focuses on the distribution and potential functions of methyltransferases in CPR, examining their involvement both as resistance-associated enzymes (MT) and as components of restriction–modification systems (hsdM).

## Materials and Methods

### Genomic data

A total of 12,552 CPR genomes were retrieved from the NCBI GenBank database on June 30, 2024 (National Center for Biotechnology Information) (35). Details regarding the analyzed genomes are provided in the Supplementary Materials (*Analyzed genomes*). All genomes were annotated *in house* using the PROKKA software.

### Reference database construction

Prior to experimental analyses involving bacterial strains expressing CPR genes, reference databases were compiled on January 5, 2024 from multiple sources, including ARG-ANNOT (24), NCBI BioProject PRJNA313047 (36), and the Comprehensive Antibiotic Resistance Database (CARD) (37). These datasets collectively represent resistance proteins associated with diverse antimicrobial agents and comprised 2,225 proteins from ARG-ANNOT, 7,187 proteins from the NCBI BioProject, and approximately 6,400 from CARD, with high methyltransferase hit incidence in a previous study (12).

From these collections, 143 proteins conferring resistance to aminoglycosides or macrolide–lincosamide–streptogramin (MLS) agents were retained, corresponding to 16S or 23S rRNA methyltransferases.

In parallel, restriction enzyme sequences were retrieved from the Swiss-Prot section of the UniProt database (38). After filtering, this dataset comprised 142 proteins belonging to type I–IV endonucleases, including 14 methyltransferases associated with restriction–modification (R–M) systems available at the same initial acquisition time (January 2024).

All databases were analyzed using the NCBI Conserved Domain Database (CDD) (39) to identify the most frequent functional domain clusters. Domain accession frequencies were visualized using scatterplots generated in RStudio (version 4.3.3) with the ggplot2 package, based on Excel files containing query sequences and corresponding domain annotations.

### Alignment of CPR proteins against reference databases

*In silico* analyses were initiated by concatenating the 12,552 CPR genomes (FAA files converted to FASTA format). Concatenated proteomes were aligned separately against the methyltransferase and endonuclease reference databases using BLASTp with an E-value cutoff of 1 × 10⁻⁴ (40). All alignments were performed locally.

Resulting hit sequences were extracted and further analyzed using CDD (39) and further filtered according to minimum alignment length thresholds defined for each cluster. A schematic overview of the alignment and filtering workflow is provided in Fig. 1. Based on these results, heatmaps illustrating the distribution of CPR methyltransferases and endonucleases were generated in RStudio (version 4.3.3) using the tidyr package for data reshaping and ggplot2 for visualization. In addition, a network—with source nodes as the CPR phyla connected to their functional enzyme targets (methyltransferases and restriction enzymes) through the functional sequences as edges—was constructed in RStudio (version 4.3.3) using the tidygraph package for network structuring and ggraph for visualization. Finally, functional CPR enzyme domain clusters were visualized in a comparative network alongside bacterial clusters to highlight both shared and distinct clusters between CPR and bacterial sequences. Network visualization and analysis were performed using Cytoscape (https://cytoscape.org/).

**Figure 1:**
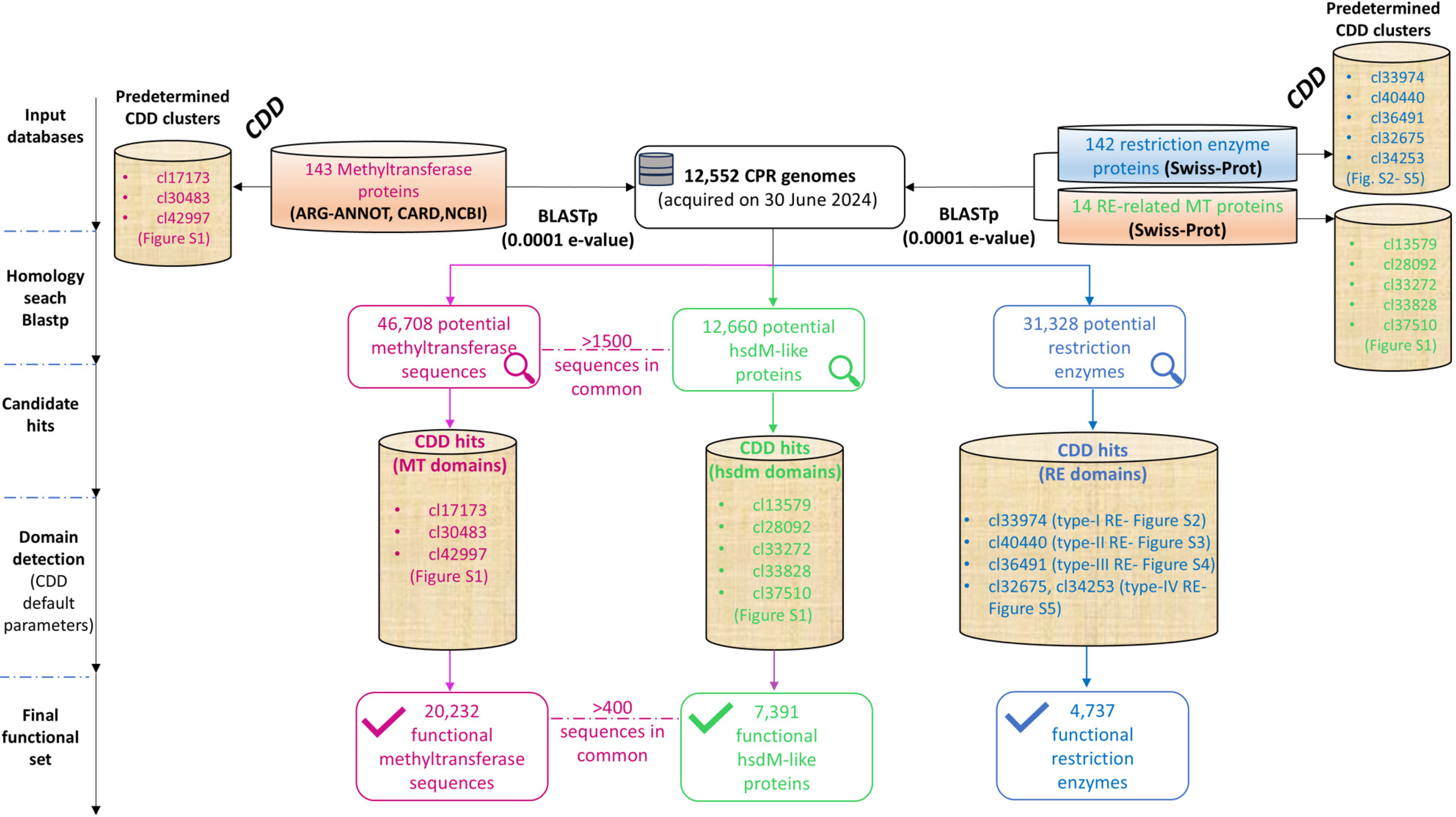
*In silico* workflow for identifying methyltransferases (MT; resistance-associated and hsdM-like proteins) and restriction endonucleases encoded in 12,552 candidate phyla radiation (CPR) genomes. Homology searches (BLASTp; E-value ≤ 1e−4) followed by conserved domain detection (CDD) were used to define candidate and functional protein sets. These functional clusters were used in order to select functional putative CPR sequence with more accuracy. A subset of sequences was shared between MT and hsdM-like groups, indicating cross-functionality or dual annotation across methyltransferase categories. Counts are shown as absolute values across enzyme classes.

Two representative CPR methyltransferase hits were selected for laboratory experiments: one corresponding to the highest-scoring BLASTp hit and one conforming to the reference domain cluster with a comparable alignment length.

### Sequence selection and validation

From the putative CPR methyltransferase dataset, two sequences were selected for *in vitro* analyses based on alignment criteria: one representing the best BLASTp hit and the other exhibiting an alignment length comparable to the reference functional domain identified by CDD. These sequences were annotated as *npmA* and *grm*, respectively. The first sequence belongs to the genome assemblies Candidate division WWE3 GCA_000993985.2 (sequence annotated as “GCA_000989435.1_ASM98943v1_00934_16S_rRNA_(guanine(1405)-N(7))-methyltransferase” and referred to as *npmA* in our study). The second sequence was extracted from Parcubacteria assembly GCA_000989435.1 (sequence annotated as “GCA_000993985.2_ASM99398v2_00181_16S_rRNA_(adenine(1408)-N(1))- methyltransferase” and referred to as *grm* in our study). Both genomes are from environmental sample origins.

Both sequences were subsequently validated through three-dimensional structural and functional predictions using the Protein Homology/Analogy Recognition Engine (Phyre2) (41).

### Cloning and transformation

The selected nucleic acid sequences were codon-optimized for *E. coli* GC content to enable *in vitro* expression. Optimized genes were synthesized by GenScript (Treubstraat 1, 2288 EG Rijswijk, Netherlands) and cloned into pET-22b(+) plasmids carrying distinct inserts. Inserts were positioned between the NdeI and NotI restriction sites.

An empty pET-22b(+) plasmid (without an insert) was used as a control. For electroporation, 1 µL of plasmid solution was added to 50 µL of electrocompetent *E. coli* BL21 (DE3) cells, with three independent attempts performed for each plasmid.

Following electroporation, bacterial mixtures were plated on LB agar (Becton, Dickinson and Company, Le Pont-de-Claix, France) supplemented with ampicillin (100 µg/mL) (EUROMEDEX, Souffelweyersheim, France) and incubated overnight at 37 °C for selection of transformed bacteria.

### BIOLOG phenotypic characterization

Phenotypic growth profiles were assessed using BIOLOG GEN III microplates according to the manufacturer’s protocol. This approach enabled visualization of growth curves for bacterial strains harboring individual plasmids compared with the control strain across different substrate wells.

GEN III plates contain 71 substrate wells (A2–H9) comprising carbon and nitrogen sources, with well A1 serving as a negative control without substrate. Wells A11–H12 assess physicochemical stress conditions (e.g., salt, antibiotics, pH variation), while well A10 serves as a positive control.

Growth curves for three bacterial strains (two test strains and one control) were monitored based on color development from reduction of a tetrazolium redox dye, reflecting substrate metabolism or resistance to physicochemical conditions.

Fresh overnight cultures were prepared in sufficient quantities to perform assays in duplicate and calibrated to a 95% transmittance. Cells were suspended in IF-A fluid, which was also used to prepare the transmittance blank. 100 µL of each prepared strain was distributed into two 96-well GEN III plates, which were then incubated in the OmniLog reader for 116 h at 37 °C.

Images were acquired at regular intervals, and instrument-specific software converted image data into numerical values to generate growth curves. Data analysis was performed 116 h after completion of data acquisition, when growth curves had largely reached a plateau, using OmniLog DATA Analysis software (version 1.7.1.53).

### Antimicrobial susceptibility testing

For aminoglycoside and macrolide susceptibility testing, each *E. coli* strain was grown to an OD₆₀₀ of 0.5–0.6, after which 125 µL of IPTG (isopropyl β-D-1-thiogalactopyranoside) (Euromedex, Strasbourg, France) was added to 25 mL of Luria–Bertani broth to achieve a final concentration of 0.5 mM and induce plasmid expression.

Additionally, 250 µL of methyl pyruvate (1 M stock) was added to the culture to obtain a final concentration of 10 mM, based on BIOLOG phenotypic results.

After incubation for 4 h at 37 °C, a 0.5 McFarland suspension was prepared for each culture. Suspensions were inoculated onto Mueller–Hinton agar (BioMérieux, Marcy-l’Étoile, France) using cotton swabs.

Minimum inhibitory concentrations (MICs) for amikacin, kanamycin, tobramycin, gentamicin, erythromycin, clarithromycin, and azithromycin were determined using E-test strips (BioMérieux) following EUCAST guidelines (https://www.eucast.org). Results were further verified using the broth microdilution method in 96-well plates, performed in parallel with E-tests, with assays conducted on Mueller–Hinton broth and agar, respectively. MIC values were recorded after 18–24 h of incubation.

## Results

### Definition of functional selection parameters for methyltransferases and restriction enzymes based on conserved domain analysis

We initiated our analyses with a domain cluster analysis of various database sequences, to determine the selection parameters of potentially function CPR sequences later. Following CDD analyses, for the methyltransferase family, the domain clusters cl17173 and cl42997, associated with 16S rRNA methyltransferases, and cl30483, representative of 23S rRNA methyltransferases, were selected based on domain complementarity (Fig. S1). In contrast, restriction enzyme domain clusters were more diverse and exhibited low commonality. Cluster cl33974 was selected for type I restriction enzymes (Fig. S2), while clusters cl40440 and cl36491 were selected for type II and type III restriction enzymes, respectively (Fig. S3 and S4). For type IV restriction enzymes, clusters cl32675 and cl34253 were retained (Fig. S5).

### High prevalence of putatively functional methyltransferases and lower abundance of restriction enzymes across CPR genomes

BLASTp alignments of 12,552 CPR genomes, using an E-value cutoff of 1 × 10⁻⁴, against both reference databases separately yielded 46,708 resistance-associated methyltransferase hits, 12,660 hsdM-like hits associated with restriction–modification (R–M) systems, and 31,328 putative restriction enzyme hits across CPR genomes (Fig. 1). BLASTp results for methyltransferases are seen in Table S1, and those for the restriction enzymes in Table S2. Then, the resulting hits were extracted and reanalyzed using CDD. CPR sequences were subsequently retained based on the presence of selected domain clusters (Fig. 1).

Following this filtering strategy, we identified 20,232 resistance-like methyltransferases (MTs), 7,391 hsdM-like proteins, and 4,737 restriction enzymes that were considered potentially functional across all CPR phyla (Fig. 1). Resulting sequences per phyla are shown in Table S3 and Table S4. The distribution of hits across the three protein classes was visualized as a node–edge network generated using RStudio (Fig. 2). Most functional proteins were associated with the unclassified Patescibacteria group, as well as the Microgenomates, Saccharibacteria, and Parcubacteria phyla. Notably, Saccharibacteria and Parcubacteria have previously been linked to human-associated environments, as mentioned above.

**Figure 2:**
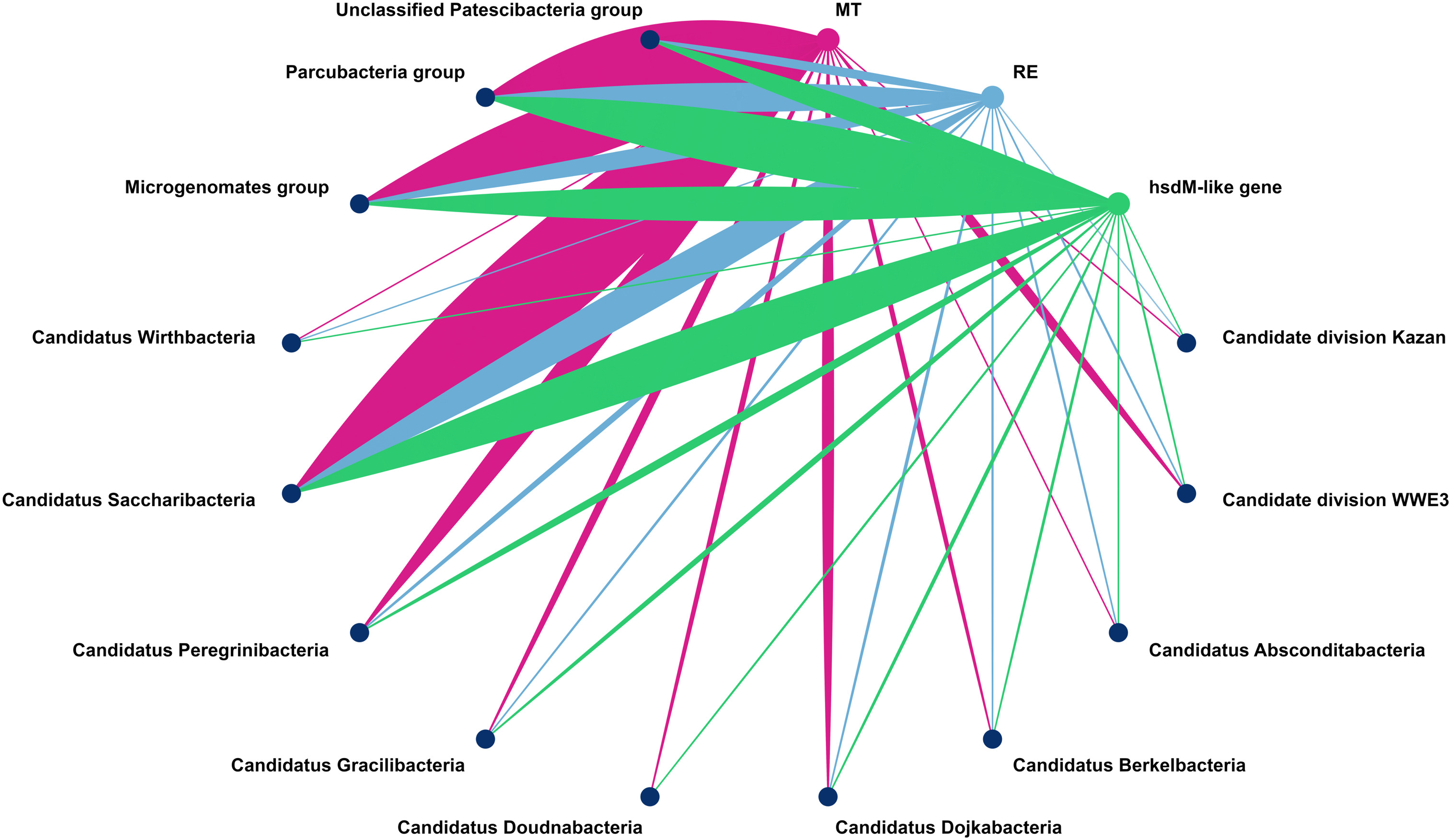
Network representation of functional Candidate Phyla Radiation (CPR) sequences across different phyla for the three protein families (i.e. MT, hsdM, RE) analyzed in this study. Navy blue nodes represent CPR lineages, and edges denote sequence associations. Edge colors indicate protein categories, functionally verified by both BLASTp and conserved domain database (CDD) analyses: pink, resistance methyltransferases; Green, hsdM methyltransferases associated with restriction–modification (R-M) systems; blue, restriction endonucleases.

To assess enzyme distribution across CPR phyla, heatmaps were generated by normalizing gene frequencies relative to the number of genomes per phylum. Heatmaps illustrated the distribution of resistance-associated MTs (Fig. S6A) and restriction enzymes and hsdM-like MTs (Fig. S6B). Gene frequencies were calculated by dividing the number of detected genes within each phylum by the total number of genomes analyzed for that phylum. Across all CPR phyla, the most prevalent resistance-associated methyltransferase gene was *erm* [13,375 hits], followed by *emt(A)* [2,418 hits], *rlmA(II)* [2,305 hits], and *cfr* [1,252 hits] methyltransferases (Fig. S6A). These genes, among others, have previously been validated as functional through CDD analyses. Despite stringent selection criteria, methyltransferase frequencies exceeded 2-3 genes and potentially reached 7 genes per phylum (as seen in Table S4). In contrast, restriction enzymes were less frequent overall (Fig. S6B), with type I restriction enzymes [3,238 hits] representing the most abundant class across CPR phyla. The relative scarcity of restriction enzymes was further reflected by the detection of 4,737 putatively functional sequences among 12,552 genomes. For hsdM-like methyltransferases, normalized frequencies ranged between 1 and 3 genes across CPR phyla, corresponding to 7,391 sequences in total (Figure S6B). All results are seen in Table S3 and TableS4. Notably, more than 1,500 sequences were classified as functional for both resistance-associated and hsdM-like methyltransferase categories, explaining the overlap observed in Fig. 1.

Altogether, our high throughput screening indicates that methyltransferase-like proteins greatly outnumber restriction enzyme-like proteins across CPR genomes.

### Extensive divergence of functional CPR sequences from known bacterial proteins

A large proportion of the identified functional CPR sequences exhibited low sequence similarity to their bacterial homologs, with most sharing less than 50% amino acid identity. Indeed, for all three enzyme classes, the majority of CPR sequences displayed only 17–30% similarity to sequences present in established functional databases (Fig. 3), indicating substantial divergence from previously characterized proteins.

**Figure 3:**
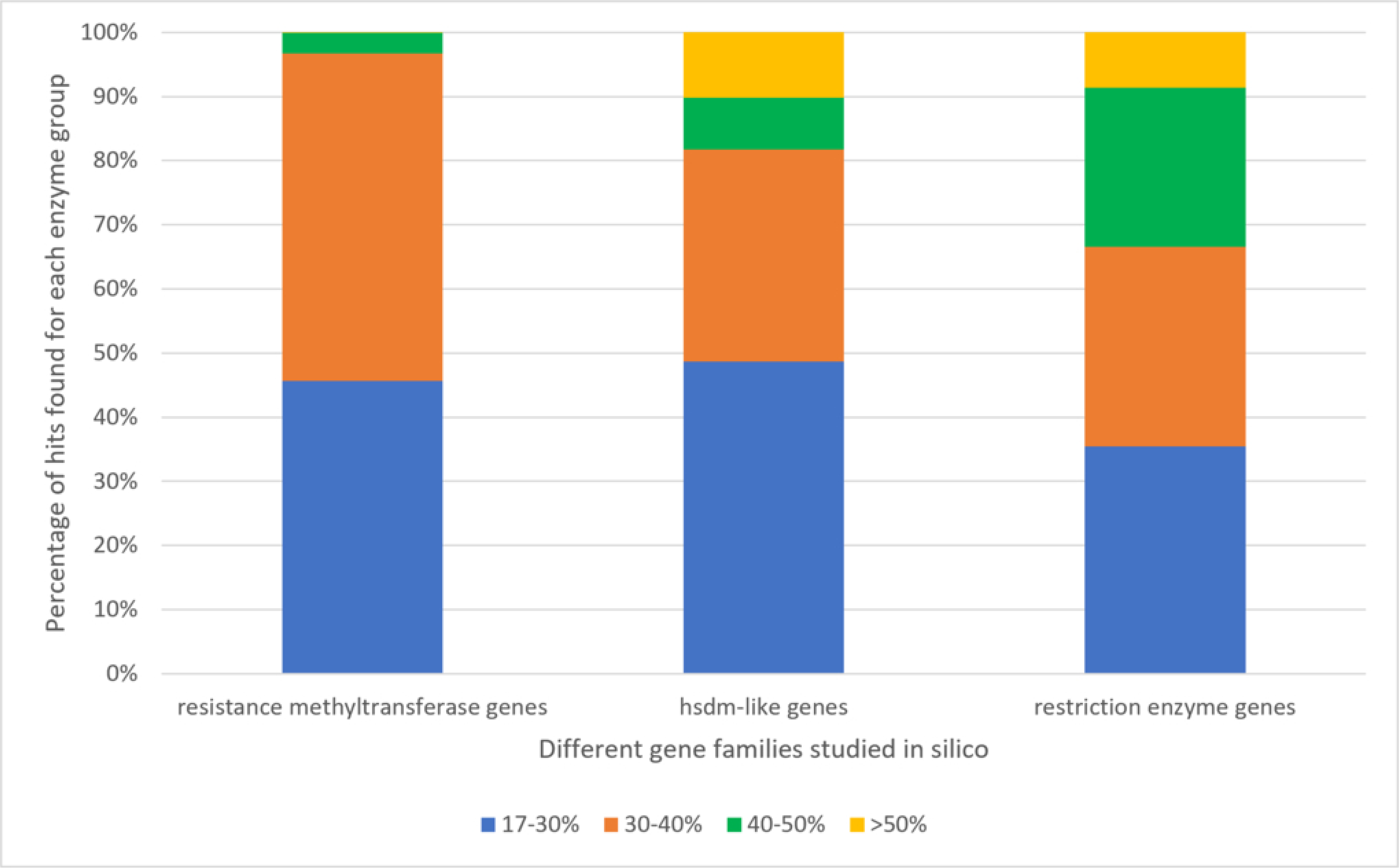
Distribution of sequence similarity to bacterial homologs across CPR methyltransferase and restriction enzyme families identified in this study. Histograms show the percentage of hits within defined similarity ranges (17–30%, 30–40%, 40–50%, and >50%) for each enzyme group. Similarity values span from 17% to 100%, indicating substantial divergence of candidate phyla radiation (CPR) protein sequences relative to bacterial counterparts.

Furthermore, only 69.5% (18,098/27,205) of the putatively functional methyltransferases were originally annotated as methyltransferases (Table S3). Consequently, more than 30% of the identified functional CPR methyltransferases had been assigned alternative annotations, further underscoring the atypical nature of CPR-associated proteins. A similar observation was reported in a previous study (12), in which 34% of antimicrobial resistance-related hits identified in CPR genomes were annotated as hypothetical proteins.

Together, the high proportion of non-canonical annotations and the low sequence similarity to known proteins highlight the remarkable divergence of CPR functional sequences from characterized bacterial proteins. These findings suggest that CPR enzymes may possess unique structural and functional features, potentially reflecting distinct evolutionary trajectories and mechanisms of action.

### CPR functional methyltransferases and restriction enzymes display greater domain diversity and functional complexity than bacterial homologs

CPR-derived functional sequences were then clustered based on conserved domain database (CDD) annotations and compared with bacterial counterparts. A total of 1,000 clusters were assigned to MT resistance proteins, with 49 clusters shared with 58 bacterial domains. CPR hsdM-like sequences formed 436 clusters, of which 18 were shared with 19 bacterial clusters. Similarly, CPR restriction enzymes generated 551 clusters compared to 165 bacterial clusters, with only 70 shared. These relationships were visualized as a node–edge network constructed in Cytoscape (Fig. 4). Overall, the network reveals a broader diversity of protein clusters in CPR compared with bacterial proteins. Notably, CPR clusters displayed a markedly different functional organization. Whereas bacterial sequences were dominated by conserved and modular restriction–modification (R–M) systems with clear separation of methyltransferase, specificity, and restriction activities, CPR sequences exhibited extensive domain mixing and multifunctionality. Methyltransferase-associated clusters frequently combined DNA, rRNA, tRNA, and protein methylation domains within single proteins, including hsdM- (cl33828 / cl33272: type I restriction-modification DNA methylase subunit involved in DNA adenine methylation), Trm- (cl34076 / cl44012: tRNA methyltransferases involved in tRNA modification), Rsm- (cl34491: rRNA methyltransferase involved in ribosomal RNA modification), and HemK-related families (cl34511: protein methyltransferase modifying release factors during translation). In addition, CPR datasets showed reduced representation of canonical restriction enzyme architectures and enrichment of helicase-associated and non-classical defense systems, including BREX-related methyltransferases (cl40397: PglX-like methyltransferase associated with phage exclusion systems) and helicase-linked domains such as DEAD-like helicases (cl28899: ATP-dependent nucleic acid helicases) and SWI2/SNF2 ATPases (cl40300: DNA/RNA remodeling ATPases), often associated with classical restriction enzyme components such as PD-(D/E)XK nucleases (cl40440: catalytic core of restriction endonucleases) and hsdR-related proteins (cl36622 / cl34711: restriction subunit of type I R–M systems with ATP-dependent nuclease activity).

**Figure 4:**
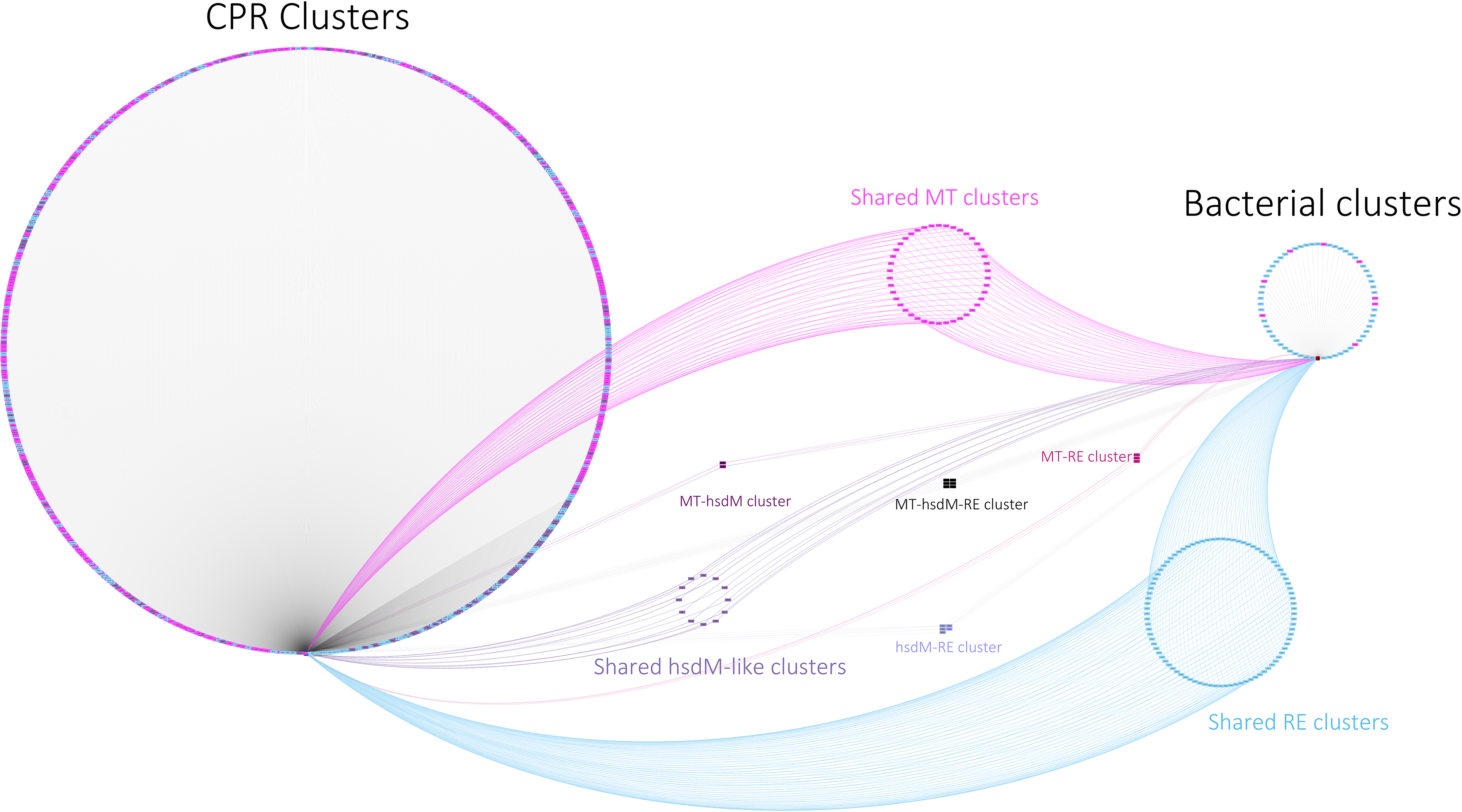
Visualization of bacterial and candidate phyla radiation (CPR) protein clusters, including shared clusters for methyltransferases (MT) in pink, hsdM-like gene encoded sequences in purple clusters, and restriction enzymes (RE) in blue clusters. Overlapping clusters highlight proteins common to multiple functional groups (MT–hsdM, MT–RE, hsdM–RE, and MT–hsdM–RE). The distribution reveals a greater diversity of CPR protein profiles among putatively functional proteins identified in this study.

Collectively, these observations indicate that CPR proteins exhibit broader and more heterogeneous functional profiles than the conserved and specialized activities observed in bacterial counterparts. This can be seen within the singularity of bacterial edge links to clusters. However, CPR sequences frequently have multiple edge links to different cluster nodes.

### Phenotypic growth curves of transformed *E. coli* strains and antimicrobial susceptibility testing

To support *in vitro* analyses, two representative sequences from the aminoglycoside methyltransferase family were selected based on highest sequence similarity to *npmA* and *grm* reference genes. These sequences were chosen for heterologous expression in *Escherichia coli* BL21 using the pET-22b(+) plasmid system. MLS-targeting genes were excluded due to the intrinsic resistance of Gram-negative bacteria to macrolides, largely driven by outer membrane impermeability and efflux-mediated exclusion (25).

Prior to *in vitro* analyses, BIOLOG GEN III microplates were used to determine optimal growth conditions for transformed *E. coli* strains. Among the tested substrates, methyl pyruvate, located in well G2, was identified as particularly relevant to this study. This observation raised the question of whether methyltransferase expression could influence methyl pyruvate metabolism.

Growth curve profiles differed markedly among the three strains, with pronounced differences observed between the control BL21 strain and strains expressing *npmA* or *grm*. In well G2, methyl pyruvate exerted an inhibitory effect on the growth of the control strain, whereas transformed strains exhibited increased tolerance, particularly the *npmA*-expressing strain. For the *grm* strain, mean values remained positive as well and clearly distinct from the negative averages recorded for the control strain. Overall, the presence of plasmids encoding *npmA*- or *grm*-like methyltransferases conferred the ability to metabolize methyl pyruvate, in contrast to the control BL21 strain, which likely lacks a functional methyltransferase involved in this process.

Antimicrobial activity was evaluated by determining minimum inhibitory concentrations (MICs) for three *E. coli* strains expressing *npmA*, *grm*, or no methyltransferase gene. MIC values, assessed using both E-test strips and 96-well broth microdilution assays, remained within a similar range compared with the control strain lacking a methyltransferase-encoding plasmid.

Bacterial growth was assessed under multiple culture conditions, including LB agar alone, LB agar supplemented with IPTG, methyl pyruvate, and both. Across all tested conditions, MIC values remained largely consistent among strains. Minor variations, typically differing by one dilution step, were observed but did not indicate a significant alteration of antimicrobial resistance patterns attributable to the expressed genes. These results are summarized in Table 1.

**Table 1:** MIC results of three *E.coli*-BL21 strains containing different modified CPR methyltransferase genes (*npmA* and *grm*), in different culture media supposedly prompting plasmid expression

| Three E.coli strains tested | MICs for different antimicrobial agents tested (µg/ml); on Mueller Hinton Agar/ Broth |  |  |  |  |  |  | Initial growth media tested |
| --- | --- | --- | --- | --- | --- | --- | --- | --- |
|  | Gentamicin<br>MIC | Tobramycin<br>MIC | Kanamycin<br>MIC | Amikacin<br>MIC | Erythromycin<br>MIC | Clarithromycin<br>MIC | Azithromycin<br>MIC |  |
| *E.coli*-BL21-*npmA* gene | 1 | 0.75 | 1.5 | 1 | 62.5 | 1000 | 125 | LB medium |
| *E.coli*-BL21-*grm* gene | 1 | 0.75 | 2 | 1.5 | 62.5 | 1000 | 125 |  |
| *E.coli*-BL21-control | 1 | 0.75 | 1.5 | 1.5 | 62.5 | 1000 | 125 |  |
| *E.coli*-BL21-*npmA* gene | 1 | 0.75 | 2 | 2 | 62.5 | 1000 | 125 | LB+ Methyl Pyruvate medium |
| *E.coli*-BL21-*grm* gene | 1 | 0.75 | 2 | 1.5 | 62.5 | 1000 | 125 |  |
| *E.coli*-BL21-control | 0.75 | 0.75 | 2 | 1.5 | 62.5 | 1000 | 125 |  |
| *E.coli*-BL21-*npmA* gene | 1.5 | 0.5 | 1.5 | 2 | 125 | 1000 | 125 | LB+ IPTG medium |
| *E.coli*-BL21-*grm* gene | 1 | 0.75 | 1.5 | 1.5 | 62.5 | 500 | 62.5 |  |
| *E.coli*-BL21-control | 1 | 0.75 | 1.5 | 1.5 | 62.5 | 500 | 125 |  |
| *E.coli*-BL21-*npmA* gene | 1.5 | 0.5 | 1.5 | 2 | 125 | 1000 | 125 | LB+ IPTG+ Methyl Pyruvate medium |
| *E.coli*-BL21-*grm* gene | 1 | 0.75 | 1.5 | 1.5 | 62.5 | 500 | 62.5 |  |
| *E.coli*-BL21-control | 1 | 0.75 | 1.5 | 1.5 | 62.5 | 500 | 62.5 |  |

## Discussion

Many aspects of CPR biology remain poorly understood, particularly in relation to the extensive loss of metabolic pathways observed in their genomes (3). The inability of CPR organisms to synthesize essential metabolites, including nucleotides and amino acids, underscores the need to better understand how these bacteria persist and adapt to their ecological niches.

Numerous resistance-like profiles have been identified within CPR genomes, including genes associated with resistance to glycopeptides, aminoglycosides, beta-lactams, and MLS antibiotics (12). Among these, CPR-encoded beta-lactamases have been particularly well studied (12,21,22). These enzymes were shown to exhibit functional properties extending beyond antibiotic degradation, including RNase activity, leading to their reclassification as metallo-hydrolases (22). In this context, our study focused on aminoglycoside- and MLS-associated methyltransferases, which were found to be highly abundant across CPR genomes.

Consistent with previous observations, methyltransferases were widely distributed among CPR phyla, with frequencies ranging from one to three copies per genome and, in some cases, exceeding these values. The *erm* gene represented the most prevalent resistance-associated methyltransferase (Fig. S6A), suggesting a potential link to 23S rRNA methylation and antimicrobial resistance. However, it is important to note that our stringent selection criteria likely excluded a substantial number of functional CPR sequences. Moreover, CPR-encoded methyltransferases exhibited relatively low sequence similarity to bacterial counterparts (in majority ranging from 17-30 %, globally below 50%), as previously reported (12), a pattern that was also observed in our dataset (Fig. 3 and Table S1, TableS2).

Despite the widespread distribution of methyltransferases across CPR phyla (Figures S6A and S6B; Fig. 2; Table S1), functional assays did not reveal a clear antimicrobial resistance phenotype in transformed *E. coli* strains. MIC values for strains expressing the *npmA*-like or *grm*-like genes remained comparable to those of the control strains (Table 1), raising questions regarding the primary role of these genes in antimicrobial resistance. Similarly, the elevated MICs observed for macrolides in transformed *E. coli* strains are unlikely to result from expression of the cloned genes, as control strains exhibited comparable MIC values, consistent with the intrinsic reduced susceptibility of Gram-negative bacteria to macrolides.

One possible explanation is related to the physiology of CPR organisms. CPR bacteria are predicted to rely primarily on fermentative metabolism, which may limit the activity of aminoglycosides. Indeed, aminoglycosides require an active electron transport chain for cellular uptake (43–45), a process that is likely absent or severely reduced in CPR organisms.

Consequently, CPR bacteria may possess an intrinsic resistance to aminoglycosides, suggesting that the identified 16S rRNA methyltransferase-like genes in CPR genomes are unlikely to have evolved primarily as aminoglycoside resistance determinants.

Furthermore, the CPR genome harboring the *npmA*-like sequence encodes four methyltransferases but no detectable R–M system enzymes (Table S4). In contrast, the genome carrying the *grm*-like sequence encodes two *hsdM*-like proteins, three additional methyltransferases, and one restriction enzyme (Table S4). These genomic contexts suggest that the tested genes may participate in R–M-related processes or possess methyltransferase functions that were not assessed in the present study.

Collectively, these findings indicate that CPR methyltransferases may play roles extending beyond classical antimicrobial resistance and may instead contribute to alternative cellular or defense-related functions.

One plausible explanation is the involvement of methyltransferases in self-DNA methylation as part of restriction–modification (R–M) systems, thereby protecting genomic DNA from endonuclease-mediated cleavage (23). This hypothesis is supported by the identification of numerous hsdM-like methyltransferases and restriction enzymes within CPR genomes.

Additionally, more than 1,500 sequences in our dataset were classified as potentially functional for both resistance-associated and hsdM-like methyltransferase categories, indicating possible functional overlap (Fig. 1). Notably, as illustrated in Fig. 4, CPR proteins exhibited broader and more heterogeneous functional clusters than their bacterial counterparts. While bacterial sequences were typically associated with a single functional cluster per domain, CPR sequences frequently contained multiple functional domains within a single protein, including domains involved in DNA, RNA, and protein methylation (Fig. 4). Moreover, several identified methyltransferases contained the cl40397 PglX methyltransferase domain, which is associated with phage exclusion systems and the inhibition of phage replication. Since this domain is not part of canonical R–M systems, its presence further highlights the functional versatility of CPR proteins. Similarly, CPR restriction enzyme-like sequences exhibited functional associations extending beyond those of classical restriction enzymes, showing increased linkage to helicases and alternative defense systems. This extensive multifunctionality clearly distinguishes CPR R–M-associated proteins from their canonical bacterial counterparts. Collectively, these findings suggest that CPR proteins have evolved multifunctional architectures, a feature that may be particularly advantageous in organisms with highly reduced genomes.

Globally, the abundance of restriction enzymes further supports this interpretation. We identified 4,737 putatively functional restriction enzymes, with type I enzymes being the most prevalent class [3, 278 hits] across CPR phyla. Most restriction enzymes belonged to type I–III methyl-dependent systems, followed by modification-dependent restriction enzymes (23,26). It is likely that our stringent filtering criteria underestimated the true abundance of functional endonucleases. Given the compact nature of CPR genomes, it is also possible that nuclease activity is embedded within multifunctional enzymes, such as CPR beta-lactamases, which have been reported to exhibit RNase and both endonuclease and exonuclease activities (12,21,27). The combined prevalence of restriction enzymes and beta-lactamases may partially offset the higher abundance of methyltransferases in CPR genomes. Thus, this distribution, consistent with the reduced genome size typical of CPR organisms, suggests a potential functional balancing in which restriction enzymes and beta-lactamases collectively approach the abundance of methyltransferases. The presence of restriction enzymes, together with beta-lactamases, may provide CPR organisms with a mechanism for nucleotide acquisition through the degradation of exogenous DNA and RNA, while methyltransferases protect self-DNA from cleavage (Fig. 5).

**Figure 5:**
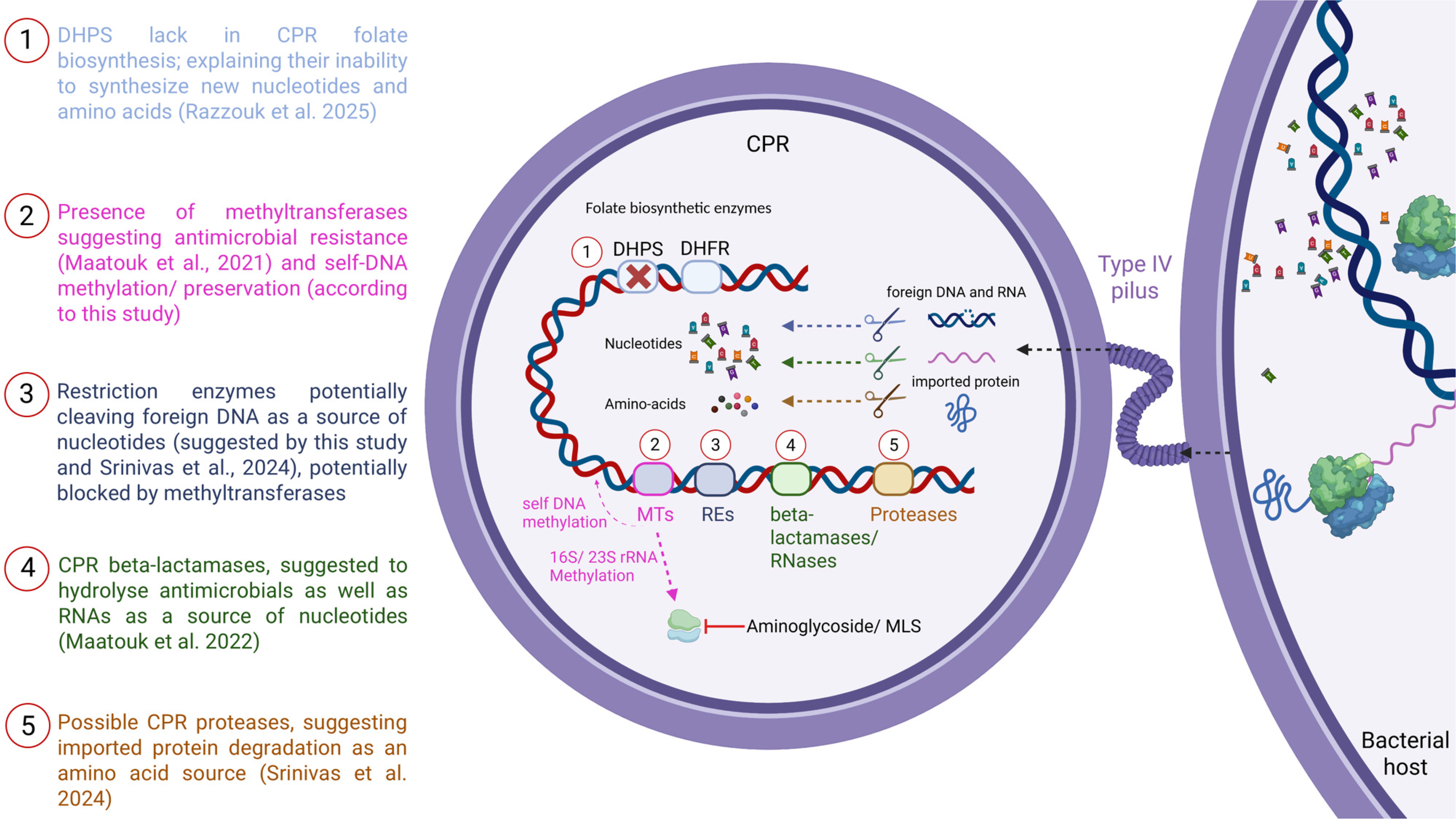
Conceptual model of self-preservation strategies in candidate phyla radiation (CPR) cells mediated by DNA methylation and macromolecule scavenging. CPR genomes lack key enzymes of the folate biosynthetic pathway (e.g., dihydropteroate synthase [DHPS]), limiting de novo nucleotide and amino acid synthesis. Restriction enzymes (RE) are proposed to cleave foreign DNA, providing nucleotides, while beta-lactamase/RNase-like proteins and proteases contribute to RNA and protein degradation, respectively, supporting nucleotide and amino acid acquisition. Methyltransferases (MTs) are implicated both in antimicrobial resistance and in protection of self-DNA via modification in restriction–modification (R–M) systems. Cleavage activities are illustrated by color-coded scissors corresponding to each functional category. This figure was created in BioRender: Bittar, F. (2026) : https://BioRender.com/l4ctyxe

Such a strategy would be particularly advantageous for organisms lacking de novo nucleotide biosynthesis pathways (28,29). Supporting this model, CPR genomes have been shown to contain numerous introns encoding endonucleases (30), further reinforcing the relevance of R–M systems in CPR biology, and the widespread distribution of endonucleases in these cells.

Taken together, our findings suggest that CPR-encoded methyltransferases and restriction enzymes form coordinated systems that support genome protection and metabolic compensation, rather than serving primarily as classical antimicrobial resistance determinants. This interpretation is consistent with recent observations linking CPR intrinsic resistance to sulfonamides with the absence of dihydropteroate synthase (DHPS), a key enzyme in folate biosynthesis (31). Additional compensatory mechanisms, including proteases (28,32), multifunctional beta-lactamases (21,22), and potential acquisition of viral DNA as a nucleotide source (33), may further contribute to CPR survival.

A schematic summary of these proposed mechanisms is presented in Fig. 5 illustrating how CPR organisms may compensate for metabolic deficiencies through coordinated cleavage- and methylation-based strategies.

## Conclusion

This study highlights the extensive presence of methyltransferases encoded by CPR genomes and supports the hypothesis that these enzymes serve multifunctional roles. Although *in silico* analyses indicate a potential association with antimicrobial resistance, our limited *in vitro* assays did not reveal a corresponding resistance phenotype, suggesting alternative biological functions.

One possibility is that CPR-encoded methyltransferases exhibit structural or functional specificity that limits their activity in heterologous bacterial systems. Alternatively, methyltransferases may primarily function within CPR-specific contexts, contributing to self-DNA protection and coordination with restriction enzymes rather than, or in addition to conferring resistance to antibiotics. Following our massive genome analyses, we proposed an *in silico* scenario (Fig. 5) highlighting the importance of methyltransferases and their possible involvement in both resistance and R-M system within the peculiar CPR physiology.

Notably, type I–III restriction enzymes were the most abundant classes identified, followed by type IV endonucleases. Type IV systems, which preferentially target methylated DNA (34), raise intriguing questions regarding their potential involvement in CPR genome streamlining and evolutionary adaptation. Such systems may contribute to genome reduction, defense against foreign DNA, and metabolic recycling.

Overall, further investigation of CPR restriction–modification systems is warranted to elucidate their functional roles, evolutionary significance, and contribution to CPR survival strategies.

Continued exploration of CPR genomes will be essential to refine our understanding of these enigmatic organisms and the unconventional biological systems they employ.

## Acknowledgment

This work was supported by a grant from the French Government managed by the National Research Agency under the “Investissements d’avenir (Investments for the Future)” programme with the reference ANR-10-IAHU-03 (Méditerranée Infection), by the Contrat Plan Etat-Région and the European funding FEDER IHUPERF.

## Declaration of generative AI and AI-assisted technologies in the writing process

During the preparation of this manuscript the authors used ChatGPT to improve the grammar of some sentences. After using this service, the authors reviewed and edited the content as needed and took full responsibility for the content of the publication.

## Transparency declaration

The authors declare that they have no competing interests.

## Authors’ contributions

R.R. and F.B. wrote the manuscript.

J.-M.R. and F.B. designed the study and revised the manuscript.

R.R. and M.B. performed the bioinformatic analyses.

R.R., L.H., M.M., and L.P. performed microbiological analyses. All authors have read and approved the final manuscript.

## Associated Data

### Supplementary Materials

#### Analyzed genomes

Within this study we relied on different CPR phyla depending on what was made available in the NCBI dataset. A total 58 genome assemblies were collected for “*Candidatus* Absconditabacteria”, 109 for “*Candidatus* Berkelbacteria”, 175 for “*Candidatus* Dojkabacteria”, 86 for “*Candidatus* Doudnabacteria”, 284 for “*Candidatus* Gracilibacteria”, 11 for the “Candidate division Kazan-3B-28”, 2,415 for the “Microgenomates group”, 4,234 for the “Parcubacteria group”, 389 for the “*Candidatus* Peregrinibacteria”, 2,661 for the “*Candidatus* Saccharibacteria”, 1,953 for the “Unclassified Patescibacteria group”, 3 for the “*Candidatus* Wirthbacteria”, and 174 genomes for the “Candidate Division WWE3” known as “Katanobacteria”.

**Table S1:** Comprehensive BLASTp results for MT and hsdM as putative methyltransferases found within 12,552 CPR genomes. All values are reported as raw counts and associated annotations as described in Materials and Methods.

**Table S2:** Comprehensive BLASTp results for RE as putative methyltransferases found within 12,552 CPR genomes. All values are reported as raw counts and associated annotations as described in Materials and Methods.

**Table S3: All functional enzymes (MT, hsdM, RE) identified in the analyzed CPR genomes, organized by phylum and by genome according to the annotated CPR sequences.**

**Table S4: Recap table, representing the number of functional CPR sequences within each protein family (MT, hsdM, RE) divised by phylum**

**Table S5: A listing of all clusters generated by the bacterial methyltransferases (MT, and hsdM) Table S6: A listing of all clusters generated by the bacterial restriction enzymes (type I- IV)**

**Figure S1:** Scatterplot representation of all functional domains found in our database. These domains were obtained through submission of reference aminoglycoside, macrolide, lincosamide, and streptogramin methyltransferases (from ARG-ANNOT, NCBI BioProject PRJNA313047 and CARD) to the NCBI Conserved Domain Database (CDD). On the y-axis are listed three complementary domain accessions (cl17173, cl30483, cl2997) for resistance-associated methyltransferases, with the remaining domains corresponding to methyltransferases associated with restriction enzymes. The listed clusters were used as selection parameters to narrow the interval of CPR functional methyltransferases.

**Figure S2:** Scatterplot representation of domain clusters identified in type I restriction enzymes, highlighting the “cl33974” cluster. **Figure S3:** Scatterplot representation of domain clusters identified in type II restriction enzymes, highlighting the “cl40440” cluster. **Figure S4:** Scatterplot representation of domain clusters identified in type III restriction enzymes, highlighting the “cl36491” cluster.

**Figure S5:** Scatterplot representation of domain clusters identified in type IV restriction enzymes, highlighting the “cl32675”, and “cl34253” clusters.

**Figure S6**: Heatmaps showing the distribution of methyltransferases (MT), HsdM-like proteins, and restriction enzymes (RE) across candidate phyla radiation (CPR) phyla. Data were generated from BLASTp and conserved domain detection (CDD) analyses of 12,552 CPR genomes. (A) Relative abundance of resistance-associated MTs across CPR phyla. (B) Distribution of restriction enzymes (type I–IV and total RE) and HsdM-like proteins across CPR phyla. Color intensity (blue gradient) represents normalized frequencies, calculated as the number of genes identified in each phylum divided by the total number of genomes assigned to that phylum. These distributions reflect enzyme prevalence under the applied stringent detection criteria.

## References

1. Ibrahim A, Colson P, Merhej V, Zgheib R, Maatouk M, Naud S, Bittar F, Raoult D. 2021. Rhizomal reclassification of living organisms. Int J Mol Sci 22:5643. doi: 10.3390/ijms22115643.

2. Ji Y, Zhang P, Zhou S, Gao P, Wang B, Jiang J. 2022. Widespread but poorly understood bacteria: Candidate Phyla Radiation. Microorganisms 10:2232. 10.3390/microorganisms10112232

3. Castelle CJ, Banfield JF. 2018. Major new microbial groups expand diversity and alter our understanding of the tree of life. Cell 172:1181–1197. doi: 10.1016/j.cell.2018.02.016.

4. Hug LA, Baker BJ, Anantharaman K, Brown CT, Probst AJ, Castelle CJ, Butterfield CN, Hernsdorf AW, Amano Y, Ise K, Suzuki Y, Dudek N, Relman DA, Finstad KM, Amundson R, Thomas BC, Banfield JF. 2016. A new view of the tree of life. Nat Microbiol 1:16048. doi: 10.1038/nmicrobiol.2016.48.

5. Méheust R, Burstein D, Castelle CJ, Banfield JF. 2019. The distinction of CPR bacteria from other bacteria based on protein family content. Nat Commun 10:4173. doi: 10.1038/s41467-019-12171-z.

6. Starr EP, Shi S, Blazewicz SJ, Probst AJ, Herman DJ, Firestone MK, Banfield JF. 2018. Stable isotope informed genome-resolved metagenomics reveals that Saccharibacteria utilize microbially-processed plant-derived carbon. Microbiome 6:122. doi: 10.1186/s40168-018-0499-z.

7. Anantharaman K, Brown CT, Hug LA, Sharon I, Castelle CJ, Probst AJ, Thomas BC, Singh A, Wilkins MJ, Karaoz U, Brodie EL, Williams KH, Hubbard SS, Banfield JF. 2016. Thousands of microbial genomes shed light on interconnected biogeochemical processes in an aquifer system. Nat Commun 7:13219. doi: 10.1038/ncomms13219.

8. Orsi WD, Richards TA, Francis WR. 2018. Predicted microbial secretomes and their target substrates in marine sediment. Nat Microbiol 3:32–37. doi: 10.1038/s41564-017-0047-9.

9. Dudek NK, Sun CL, Burstein D, Kantor RS, Aliaga Goltsman DS, Bik EM, Thomas BC, Banfield JF, Relman DA. 2017. Novel Microbial Diversity and Functional Potential in the Marine Mammal Oral Microbiome. Current Biology 27:3752–3762.e6. doi:10.1016/j.cub.2017.10.040

10. Danczak RE, Johnston MD, Kenah C, Slattery M, Wrighton KC, Wilkins MJ. 2017. Members of the Candidate Phyla Radiation are functionally differentiated by carbon- and nitrogen-cycling capabilities. Microbiome 5:112. doi: 10.1186/s40168-017-0331-1.

11. He X, McLean JS, Edlund A, Yooseph S, Hall AP, Liu SY, Dorrestein PC, Esquenazi E, Hunter RC, Cheng G, Nelson KE, Lux R, Shi W. 2015. Cultivation of a human-associated TM7 phylotype reveals a reduced genome and epibiotic parasitic lifestyle. Proc Natl Acad Sci U S A 112:244–249. doi: 10.1073/pnas.1419038112.

12. Maatouk M, Ibrahim A, Rolain JM, Merhej V, Bittar F. 2021. Small and equipped: the rich repertoire of antibiotic resistance genes in Candidate Phyla Radiation genomes. mSystems 6:e00898–21. 10.1128/mSystems.00898-21

13. Luef B, Frischkorn KR, Wrighton KC, Holman H-YN, Birarda G, Thomas BC, Singh A, Williams KH, Siegerist CE, Tringe SG, Downing KH, Comolli LR, Banfield JF. 2015. Diverse uncultivated ultra-small bacterial cells in groundwater. Nat Commun 6:6372. doi: 10.1038/ncomms7372.

14. Bernard C, Lannes R, Li Y, Bapteste É, Lopez P. 2020. Rich repertoire of quorum sensing protein coding sequences in CPR and DPANN associated with interspecies and interkingdom communication. mSystems 5:e00414–20. 10.1128/mSystems.00414-2

15. Naud S, Ibrahim A, Valles C, Maatouk M, Bittar F, Tidjani Alou M, Raoult D. 2022. Candidate Phyla Radiation, an underappreciated division of the human microbiome, and its impact on health and disease. Clin Microbiol Rev 35:e00140–21. doi:10.1128/cmr.00140-21.

16. Maatouk M, Rolain JM, Bittar F. 2023. Using genomics to decipher the enigmatic properties and survival adaptation of Candidate Phyla Radiation. Microorganisms 11:1231. doi:10.3390/microorganisms11051231.

17. Anantharaman K, Brown CT, Burstein D, Castelle CJ, Probst AJ, Thomas BC, Williams KH, Banfield JF. 2016. Analysis of five complete genome sequences for members of the class Peribacteria in the recently recognized Peregrinibacteria bacterial phylum. PeerJ 4:e1607. doi:10.7717/peerj.1607.

18. Perron GG, Whyte L, Turnbaugh PJ, Goordial J, Hanage WP, Dantas G, Desai MM. 2015. Functional characterization of bacteria isolated from ancient arctic soil exposes diverse resistance mechanisms to modern antibiotics. PLoS One 10 :e0069533. doi:10.1371/journal.pone.0069533.

19. McLean JS, Bor B, Kerns KA, Liu Q, To TT, Solden L, Hendrickson EL, Wrighton K, Shi W, He X. 2020. Acquisition and adaptation of ultra-small parasitic reduced genome bacteria to mammalian hosts. Cell Rep 32:107939. doi:10.1016/j.celrep.2020.107939.

20. Shaiber A, Willis AD, Delmont TO, Roux S, Chen LX, Schmid AC, Yousef M, Watson AR, Lolans K, Esen ÖC, Lee STM, Downey N, Morrison HG, Dewhirst FE, Mark Welch JL, Eren AM. 2020. Functional and genetic markers of niche partitioning among enigmatic members of the human oral microbiome. Genome Biol 21:292. doi:10.1186/s13059-020-02195-w.

21. Maatouk M, Ibrahim A, Pinault L, Armstrong N, Azza S, Rolain JM, Bittar F, Raoult D. 2022. New beta-lactamases in Candidate Phyla Radiation: owning pleiotropic enzymes is a smart paradigm for microorganisms with a reduced genome. Int J Mol Sci 23:5446. doi:10.3390/ijms23105446.

22. Maatouk M, Merhej V, Pontarotti P, Ibrahim A, Rolain JM, Bittar F. 2023. Metallo-beta-lactamase-like encoding genes in Candidate Phyla Radiation: widespread and highly divergent proteins with potential multifunctionality. Microorganisms 11:1933. doi:10.3390/microorganisms11081933.

23. Gao Q, Lu S, Wang Y, He L, Wang M, Jia R, Chen S, Zhu D, Liu M, Zhao X, Yang Q, Wu Y, Zhang S, Huang J, Mao S, Ou X, Sun D, Tian B, Cheng A. 2023. Bacterial DNA methyltransferase: a key to the epigenetic world with lessons learned from proteobacteria. Front Microbiol 14:1129437. doi:10.3389/fmicb.2023.1129437.

24. Gupta SK, Padmanabhan BR, Diene SM, Lopez-Rojas R, Kempf M, Landraud L, Rolain JM. 2014. ARG-ANNOT, a new bioinformatic tool to discover antibiotic resistance genes in bacterial genomes. Antimicrob Agents Chemother 58:212–220. doi:10.1128/AAC.01310-13.

25. Ma Y, Pirolo M, Jana B, Mebus VH, Guardabassi L. 2024. The intrinsic macrolide resistome of Escherichia coli. Antimicrob Agents Chemother 68:e00452–24. doi:10.1128/AAC.00452-24.

26. Bell RT, Sahakyan H, Makarova KS, Wolf YI, Koonin EV. 2024. CoCoNuTs are a diverse subclass of type IV restriction systems predicted to target RNA. eLife 13:RP100626. doi:10.7554/eLife.100626.1.

27. Lewis K. 2013. Platforms for antibiotic discovery. Nat Rev Drug Discov 12:371–387. doi:10.1038/nrd3975.

28. Castelle CJ, Brown CT, Anantharaman K, Probst AJ, Huang RH, Banfield JF. 2018. Biosynthetic capacity, metabolic variety and unusual biology in the CPR and DPANN radiations. Nat Rev Microbiol 16:629–645. doi: 10.1038/s41579-018-0076-2.

29. Gios E, Mosley OE, Takeuchi N, Handley KM. 2025. Genetic exchange shapes ultra-small Patescibacteria metabolic capacities in the terrestrial subsurface. mSystems 10:e00046–25. doi:10.1128/msystems.00046-25.

30. Tsurumaki M, Sato A, Saito M, Kanai A. 2024. Comprehensive analysis of insertion sequences within rRNA genes of CPR bacteria and biochemical characterization of a homing endonuclease encoded by these sequences. J Bacteriol 206:e00074–24. doi:10.1128/jb.00074-24.

31. Razzouk R, Azour A, Rolain JM, Bittar F. 2025. A silent sulfonamide-resistant microbial world: The curious case of missing DHPS in candidate phyla radiation. J Glob Antimicrob Resist 44:241–243. doi:10.1016/j.jgar.2025.06.024.

32. Srinivas P, Peterson SB, Gallagher LA, Wang Y, Mougous JD. 2024. Beyond genomics in Patescibacteria: A trove of unexplored biology packed into ultrasmall bacteria. Proc Natl Acad Sci U S A 121:e2419369121. doi:10.1073/pnas.2419369121.

33. Chen LX, Al-Shayeb B, Méheust R, Li WJ, Doudna JA, Banfield JF. 2019. Candidate Phyla Radiation Roizmanbacteria from hot springs have novel and unexpectedly abundant CRISPR-Cas systems. Front Microbiol 10:928. doi:10.3389/fmicb.2019.00928.

34. Sitaraman R, Leppla SH. 2012. Methylation-dependent DNA restriction in Bacillus anthracis. Gene 494:44–50. doi:10.1016/j.gene.2011.11.061.

35. National Center for Biotechnology Information. NCBI Genome. https://www.ncbi.nlm.nih.gov/datasets/genome

36. National Center for Biotechnology Information. 2016. Bacterial antimicrobial resistance reference gene database. BioProject ID 1304. https://www.ncbi.nlm.nih.gov/bioproject/313047.

37. Comprehensive Antibiotic Resistance Database. CARD: The Comprehensive Antibiotic Resistance Database. https://card.mcmaster.ca

38. UniProt Consortium. UniProt: the Universal Protein Knowledgebase. https://www.uniprot.org/

39. National Center for Biotechnology Information. Conserved Domain Database and Resources. https://www.ncbi.nlm.nih.gov/Structure/bwrpsb/bwrpsb.cgi

40. National Center for Biotechnology Information. Basic Local Alignment Search Tool (BLAST). https://blast.ncbi.nlm.nih.gov/Blast.cgi

41. Protein Homology/analogY Recognition Engine (Phyre2). Protein Homology/analogY Recognition Engine V2.0. https://www.sbg.bio.ic.ac.uk/phyre2/

42. Willemse N, Schultsz C. 2016. Distribution of type I restriction–modification systems in Streptococcus suis: an outlook. Pathogens 5:62. doi:10.3390/pathogens5040062.

43. Krause KM, Serio AW, Kane TR, Connolly LE. 2016. Aminoglycosides: an overview. Cold Spring Harb Perspect Med 6:a027029. doi:10.1101/cshperspect.a027029.

44. Mingeot-Leclercq MP, Glupczynski Y, Tulkens PM. 1999. Aminoglycosides: activity and resistance. Antimicrob Agents Chemother 43:727–737. doi:10.1128/AAC.43.4.727.

45. Bryan LE, Kowand SK, Van Den Elzen HM. 1979. Mechanism of aminoglycoside antibiotic resistance in anaerobic bacteria: Clostridium perfringens and Bacteroides fragilis. Antimicrob Agents Chemother 15:7–13. doi:10.1128/AAC.15.1.7.

